# Molecular signatures for inflammation vary across cancer types and correlate significantly with tumor stage, gender and vital status of patients

**DOI:** 10.1101/732867

**Authors:** Alexandra R. So, Jeong Min Si, David Lopez, Matteo Pellegrini

## Abstract

Cancer affects millions of individuals worldwide. One shortcoming of traditional cancer classification systems is that, even for tumors affecting a single organ, there is significant molecular heterogeneity. Precise molecular classification of tumors could be beneficial in personalizing patients’ therapy and predicting prognosis. To this end, here we propose to use molecular signatures to further refine cancer classification. Molecular signatures are collections of genes characterizing particular cell types, tissues or disease. Signatures can be used to interpret expression profiles from heterogeneous samples. Large collections of gene signatures have previously been cataloged in the MSigDB database. We have developed a web-based Signature Visualization Tool (SaVanT) to display signature scores in user-generated expression data. Here we have undertaken a systematic analysis of correlations between inflammatory signatures and cancer samples, to test whether inflammation can differentiate cancer types. Inflammatory response signatures were obtained from MsigDB and SaVanT and a signature score was computed for samples associated with 7 different cancer types. We first identified types of cancers that had high inflammation levels as measured by these signatures. The correlation between signature scores and metadata of these patients (gender, age at initial cancer diagnosis, cancer stage, and vital status) was then computed. We sought to evaluate correlations between inflammation with other clinical parameters and identified four cancer types that had statistically significant association (p-value < 0.05) with at least one clinical characteristic: pancreas adenocarcinoma (PAAD), cholangiocarcinoma (CHOL), kidney chromophobe (KICH), and uveal melanoma (UVM). These results may allow future studies to use these approaches to further refine cancer subtyping and ultimately treatment.

## Introduction

Cancer is a major public health problem with high mortality rates in the United States and worldwide and poses an enormous burden to individuals and society. Over 1.7 million newly diagnosed cancer cases and over 600,000 cancer deaths were estimated in the United States in 2018 (1). Screening for some cancers can lead to early detection (mammography for breast cancer and colonoscopy for colon cancer, for example), when local resection or definitive treatment may still be feasible (2). However, many cancers are found when there is already local invasion or even distant metastatic disease. In those cases, common treatment options include chemotherapy, locoregional therapies and radiation treatment (3). Among the issues complicating treatment options are the fact that there are many tumor types, whose response to therapy may differ depending on site of origin and cellular composition (4). Furthermore, even within the same organ, there are heterogeneous tumor types with different responses to therapies.

As a result, precise tumor classification is crucial; depending on the categorization of a tumor, the clinical course, prognosis, and treatment can vary dramatically (5). In general, there are two ways to classify cancer: the traditional histology-based method and molecular methods. The traditional method is based on observing the site of origin, degree of spread and cellular morphology, while the molecular method identifies gene expression and genetic profiles (6–8). Because tumors are heterogeneous and frequently contain abundant somatic mutations, traditional approaches for classifying tumor subtypes are often insufficient. Conventional histopathological evaluation of cancer utilizes surgical or image-guided biopsies of the primary tumor, often requiring serial samples throughout treatment course. Different parameters such as tumor size, grade, and degree of invasion along with other features such as tumor markers, atypical morphology, and regional lymph node drainage pattern are evaluated to predict tumor prognosis (9). Although this traditional approach to tumor categorization is valuable in many cases, it does not always accurately stratify patients into different treatment regimens or account for the molecular variability of cancer (10).

By contrast, molecular classification is based on the analysis of tumor genomes as well as gene expression (11). Successful molecular subdivision of tumors originating from the same tissue may result in different treatments targeting a specific tumor type, as is found in the case of ERBB2-amplified breast cancer and EGFR mutant lung carcinoma (12, 13). Furthermore, techniques such as RNA sequencing (RNAseq) can be used to obtain a profile of tumor RNA levels to study mutations, alternative splicing, and gene expression levels (14). Molecular signatures may also be utilized to inform biological interpretations. Molecular signatures are collections of genes with associated biological processes that can identify genes upregulated in specific sample subsets when compared to broader groups (15). Signatures can be composed of genes associated with specific diseases; for instance, breast cancer molecular signatures have identified subphenotypes indistinguishable by traditional histologic analysis (15, 16).

Mutations resulting in cancer may come about by a variety of sources, including inflammation. Chronic inflammation has been shown to increase cancer risk (17, 18) by causing tumor initiation, promotion, and metastatic progression (19). Many environmental causes of cancer are related to chronic inflammation. As many as 20% of cancers are associated with chronic infection, 30% with tobacco smoking and inhaled pollutants such as asbestos, and 35% with dietary factors (20–23). Chronic disease exposing patients to inflammation are also associated with increased cancer risk. Inflammatory bowel disease (i.e. ulcerative colitis and Crohn’s disease) is associated with an increased risk of colon adenocarcinoma (24), chronic pancreatitis is a significant risk factor for pancreatic cancer (25), and chronic gastritis secondary to *Helicobacter pylori* infection is associated with the majority of gastric cancer cases (26).

Several large consortia, such as The Cancer Genome Atlas (TCGA), provide tools and data to study the molecular basis of cancer (11, 27). The purpose of our study is to understand molecular patterns related to inflammation. Although TCGA started out by collecting only three cancer types – glioblastoma multiforme, lung, and ovarian cancers – it expanded rapidly; by 2014, genomic characterization and sequence analysis had been completed for 33 cancer types with data for over 12,000 individuals (27).

Signature visualization of individual samples allows identification of patient subcategories a priori on the basis of well-defined molecular signatures (15). As such, data from TCGA could potentially be utilized to obtain and evaluate molecular signatures. To overcome limitations of existing tools to evaluate molecular signatures, the Signature Visualization Tool (SaVanT) was previously developed as a web-based tool to visualize signatures in user-generated expression profiles (15). SaVanT has been utilized to distinguish signature scores in patients with various conditions such as infections and leukemia, providing insight into immune response of various skin diseases (15). By visualizing molecular signatures, SaVanT allows users to efficiently leverage pre-existing biological knowledge (such as from sources like TCGA) to interpret transcriptomic experiments (15).

To our knowledge, no systematic study utilizing gene signatures to evaluate tumor inflammation in TCGA has been carried out. Therefore, in this study, we aimed to use SaVanT to evaluate molecular signatures obtained from TCGA to evaluate the relationship between clinical status and inflammatory responses across multiple cancer types.

## Materials and Methods

### Data collection

In order to examine the role of inflammation in different cancer types, we sought to utilize a large-scale, systematically-processed dataset. We chose to analyze data from TCGA. All of the data has been processed with a uniform analysis pipeline, allowing for robust comparison across samples and tumor types. Gene expression data was retrieved from TCGA using their web-accessible data portal. In order to ensure that the data was normalized, we used their Harmonized Data Portal to access the data and did not include any datasets processed independently from the harmonized data. For all TCGA projects, we downloaded RNAseq data as normalized counts for all patients. Individual files (one per patient) were combined into a single matrix per primary site. In order to focus the evaluation of our methods, seven different tumor primary types were chosen to be utilized for analysis with clinical metadata – pancreatic adenocarcinoma (PAAD), glioblastoma multiforme (GBM), cholangiocarcinoma (CHOL), kidney renal papillary cell carcinoma (KIRP), kidney chromophobe (KICH), adrenocortical carcinoma (ACC), and uveal melanoma (UVM). We chose these seven out of the thirty-four tumor primary sites on TCGA, to obtain a range of inflammatory states estimated based on our analyses described below. Four types of data were retrieved for each sample: gender, age at initial cancer diagnosis, cancer stage, and vital status.

### Quality control

All data retrieved from TCGA was inspected for consistency by making sure that all profiles contained the same number of genes and that patient data was not redundant or duplicated. Furthermore, the distribution of normalized counts was analyzed at both the patient level as well as primary tumor site to identify any outliers or issues with normalization. Patient-level data was averaged to determine a single value for all genes per primary site.

### Comparison to molecular signatures

Molecular signatures were taken from the repository MSigDB (28) and SaVanT (15). Several studies and efforts have sought to identify genes involved in inflammatory pathways (28–31). Many of these projects have produced inflammatory signatures, which catalog the genes most important in several inflammatory states. In order to determine the role of inflammation across the 7 cancer types in our analysis, we utilized the ‘hallmark inflammation’ signature from MSigDB (28), a repository of molecular signatures. This signature includes 200 genes associated with acute and chronic inflammation responses, as well as elements of the TGF-β signaling cascade.

### Data and Statistical Analysis

Patient metadata was compared with the corresponding hallmark inflammatory response. Metadata and corresponding statistical tests were utilized as follows – Age (Pearson correlation), Stage (Anova Single Value), Sex (Anova Single Value), and Vital Status (Anova Single Value). Statistical significance was set at p < 0.05. Utilizing this data, box-whisker plots were created for metadata and inflammatory response correlations that had significant P-values.

## Results

### Tumor Types

Hierarchical clustering was performed to group cancer subtypes by inflammatory signature scores, and the three subgroups were determined by the dendrogram structure resulting from the hierarchical clustering (Figure 1). Of the tumor types evaluated in our study, we found that tumors in areas exposed to airways or gastrointestinal tracts, including pancreatic, lung and esophageal cancer, tended to be more inflammatory.

Based on these results we selected seven tumor types with varying levels of inflammation for further analysis (Figure 2). Of the 7 tumors chosen for analysis, the levels of inflammation in descending order are summarized in table 3.

**Figure 1:**
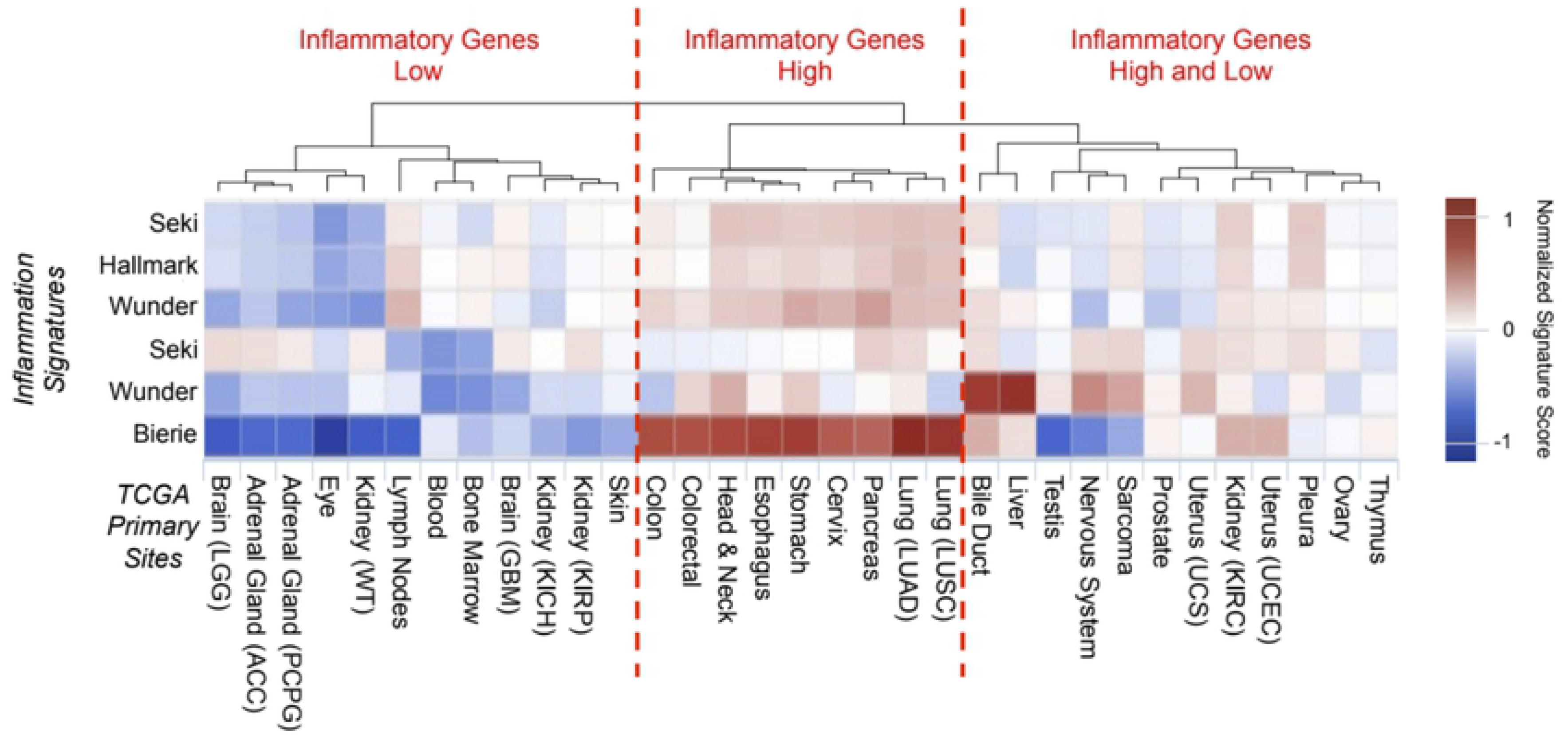
Using 6 inflammation signatures, we clustered cancer types into low, high and mixed inflammatory response.

**Figure 2:**
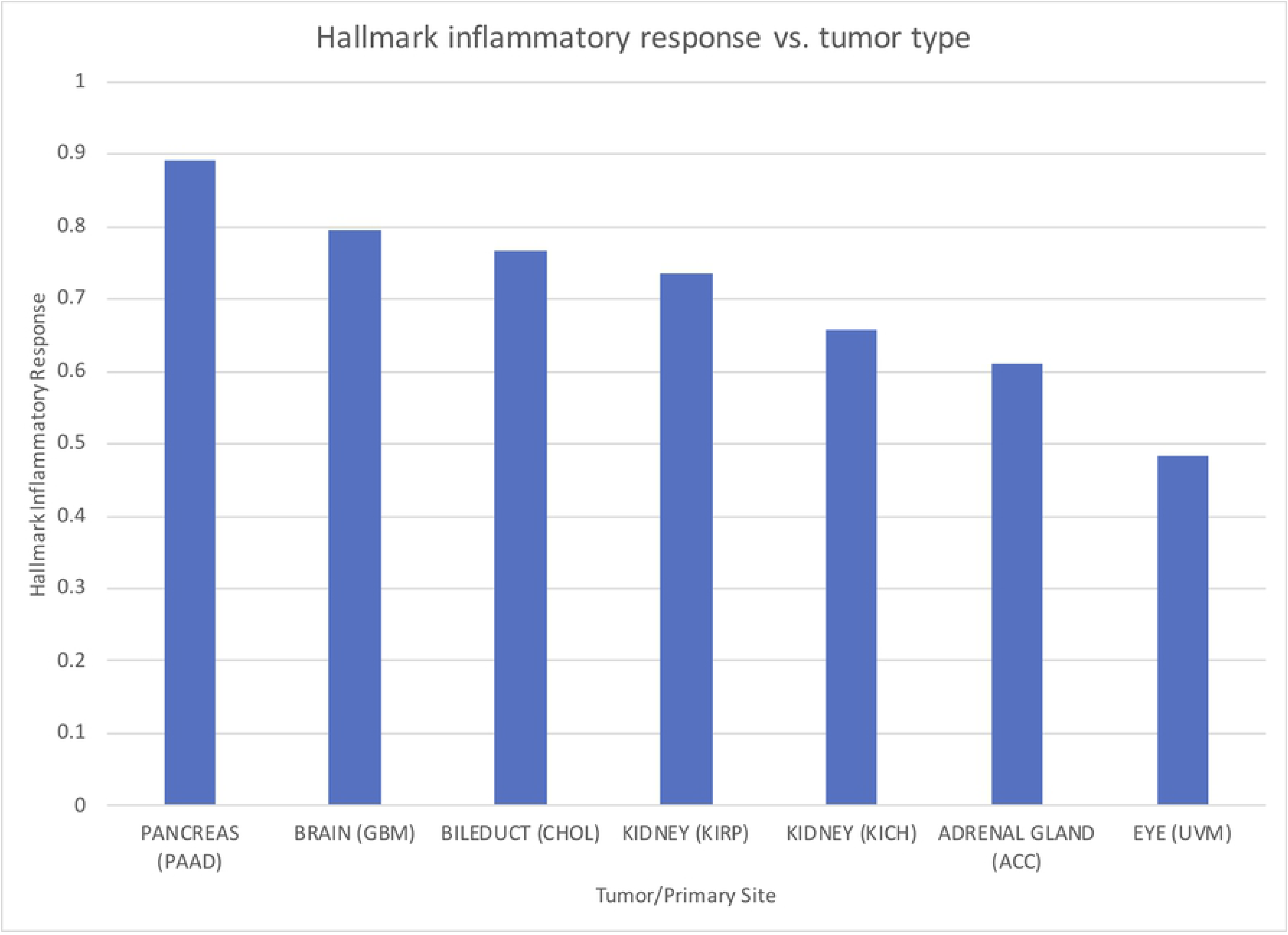
The seven chosen cancer types display heterogeneity in hallmark inflammatory response signatures.

### Correlation of Metadata with Hallmark Inflammatory Response Signature Correlation

Based on a statistical analysis of the metadata and its association with the hallmark inflammatory signature score, we were able to determine significant associations between inflammation and other clinical characteristics across specific primary cancers. Out of the 7 primary cancers, 4 had statistically significant association (p-value < 0.05) with at least one of the clinical values that we tested (sex, vital status, age at initial diagnosis, and tumor stage) as follows: pancreatic adenocarcinoma (PAAD), cholangiocarcinoma (CHOL), kidney chromophobe (KICH), and uveal melanoma (UVM).

### Pancreas Adenocarcinoma (PAAD)

After associating the metadata of 182 patients with the hallmark inflammatory response signature in pancreatic adenocarcinoma samples, we found a significant association of the hallmark inflammatory response signature with sex (p = 0.0313) and tumor stage (p = 0.0054) (Figure 3). Of all tumor stages, stage II showed the highest level of the Hallmark Inflammatory Response, and stage I the lowest. There was a slightly higher level of inflammation in females than males.

**Figure 3:**
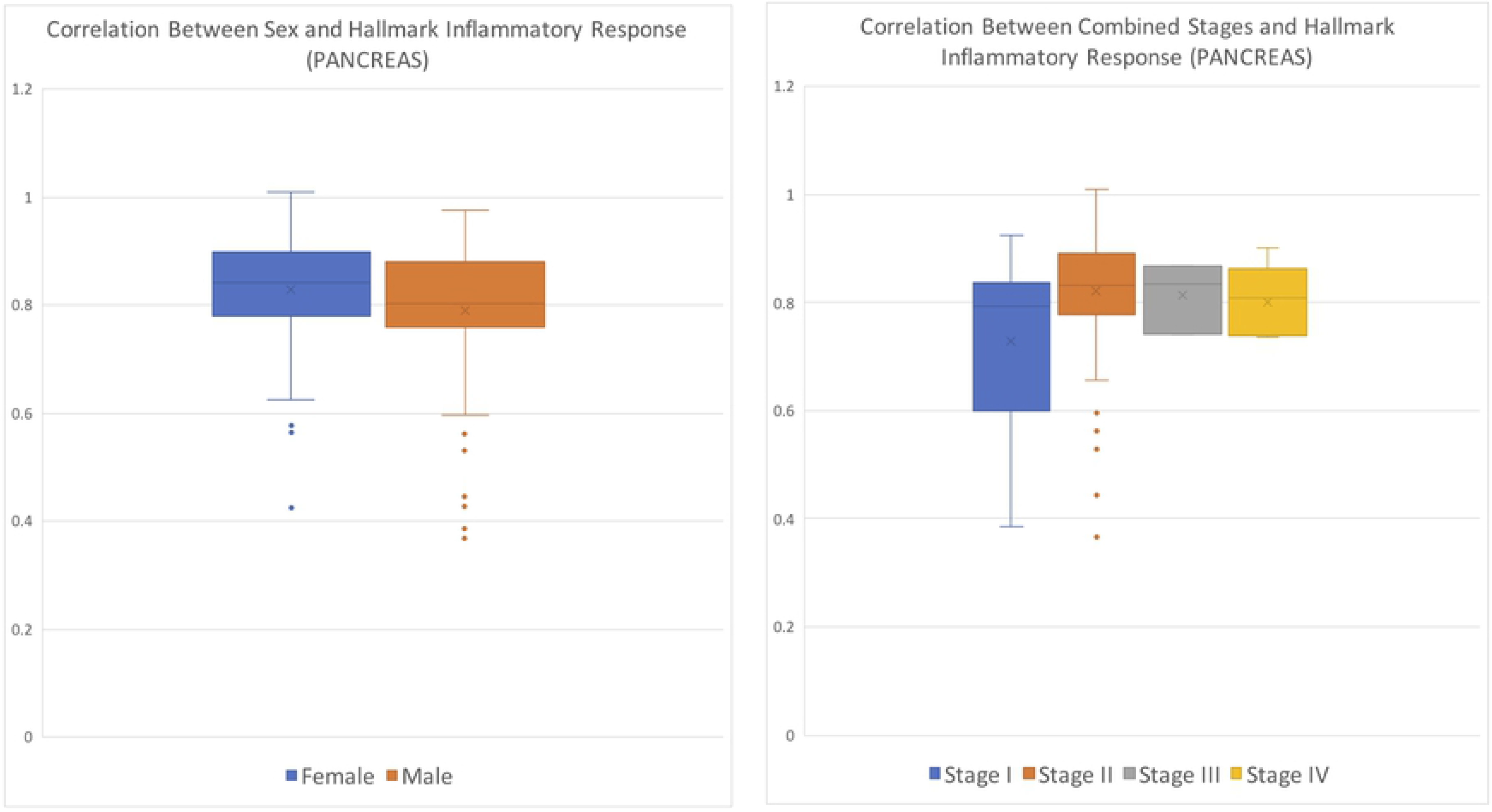
Pancreatic adenocarcinoma correlation of cancer stage with hallmark inflammatory response, p-value of 0.0054. Pancreatic adenocarcinoma correlation of sex with hallmark inflammatory response, p-value of 0.0313.

### Cholangiocarcinoma (CHOL)

We found a significant p-value of 0.0496 (Figure 4) for the association of cholangiocarcinoma inflammatory score in 45 patients with the vital status of patients, a clinical data element categorized under diagnosis on TCGA. Patients who were alive at the time of diagnosis had higher levels of inflammation than patients whose biopsy was collected post mortem.

**Figure 4:**
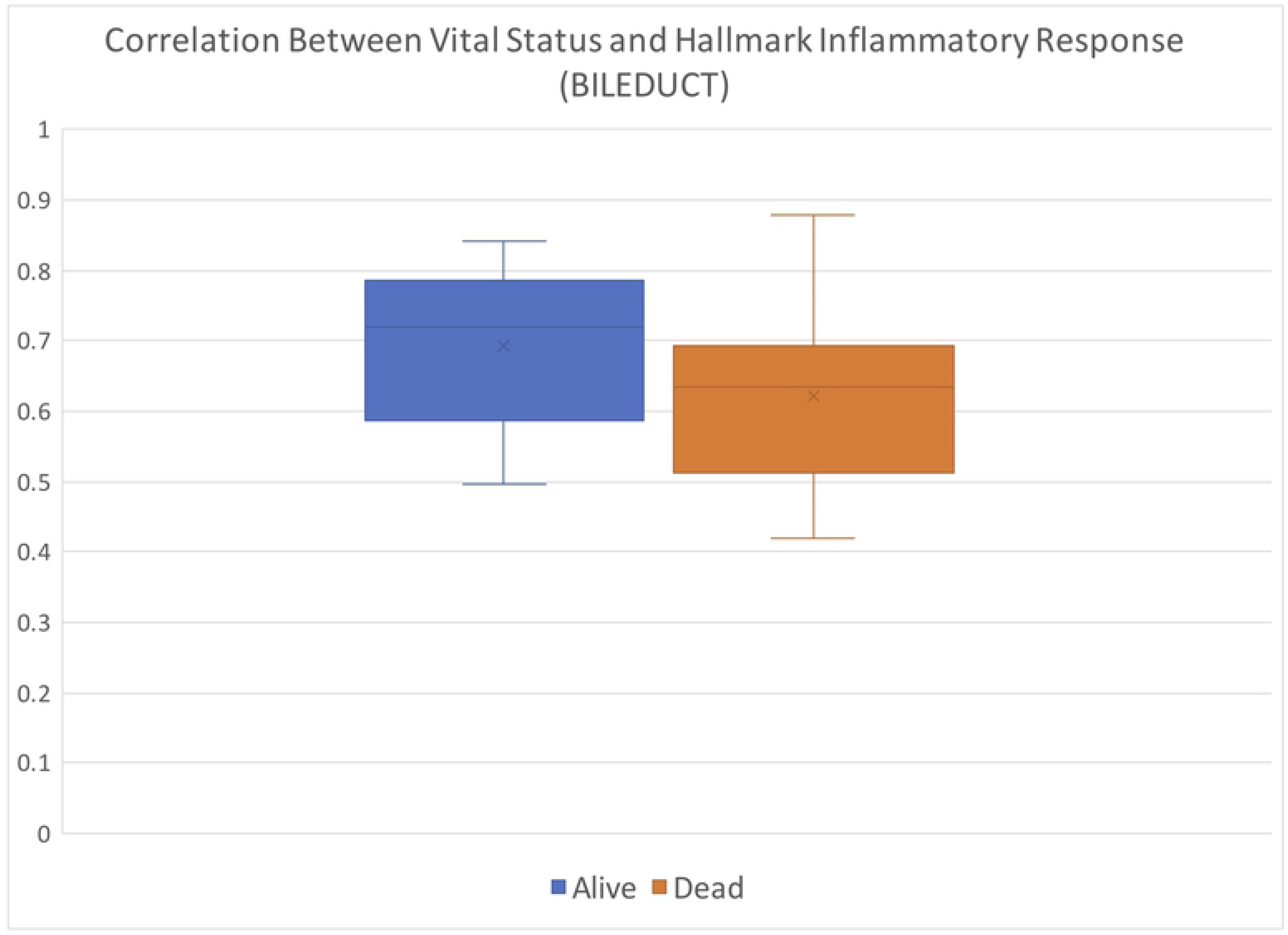
Cholangiocarcinoma correlation of vital status with hallmark inflammatory response, p-value of 0.0496.

### Kidney Chromophobe (KICH)

Using the signatures and metadata of 89 patients, we found a significant p-value = 0.0172 (Figure 5) between tumor stage and the hallmark inflammatory response. Stage IV tumors showed the highest levels of inflammation, compared to the other 3 stages.

**Figure 5:**
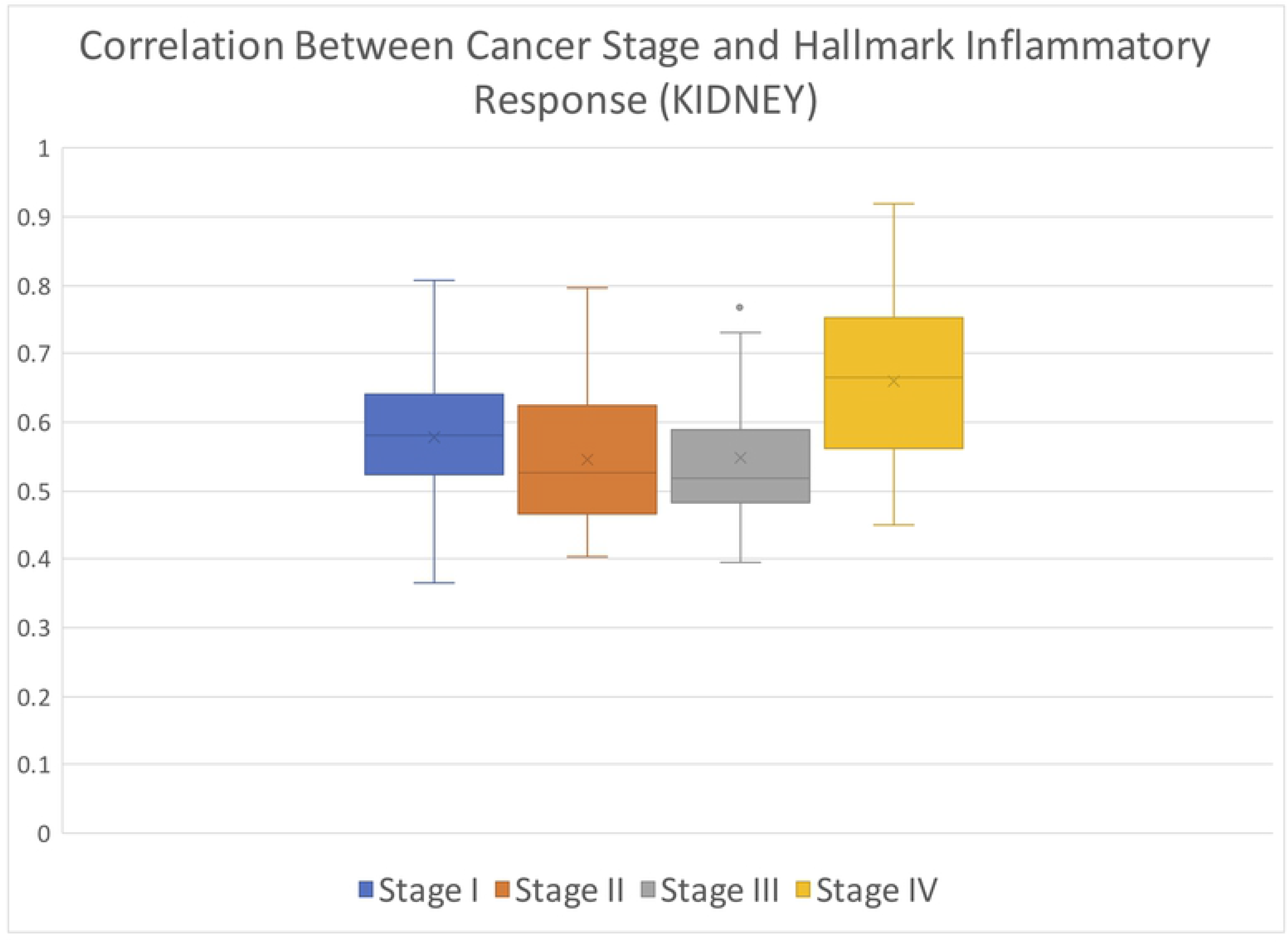
Kidney chromophobe correlation of cancer stage with hallmark inflammatory response, p-value of 0.0172.

### Uveal Melanoma (UVM)

After associating the vital status with the hallmark inflammatory response signature correlation of 80 patients, we found a significant p-value of 0.0033 (Figure 6). Overall, samples collected post mortem had higher levels of inflammation compared to those collected from living patients.

**Figure 6:**
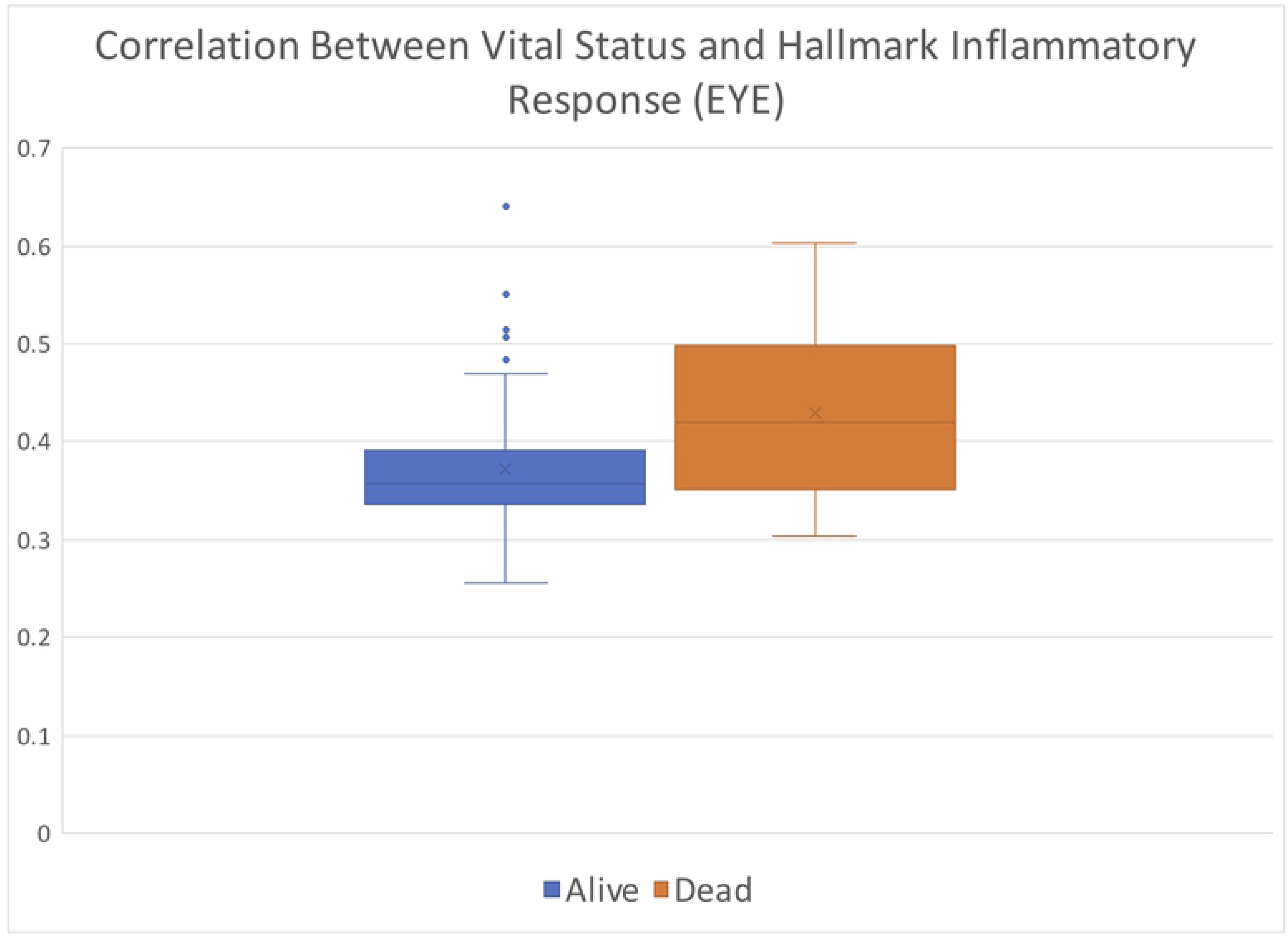
Uveal melanoma correlation of vital status with hallmark inflammatory response, p-value of 0.0033.

## Discussion

Utilizing the TCGA database allows us to leverage the systematic profiling of thousands of tumors from individuals with different types of cancer. The first aim in our study was to evaluate levels of inflammation across tumor types utilizing appropriate signature scores. We used inflammation signatures to analyze the gene expression data (“hallmark inflammation” (28–31). These signatures allow us to classify the cancer subtypes at an immunological level, which is not possible with traditional classification schemes relying on histological data. Such a classification technique allows us to examine individual pathways and signaling cascades, particularly those important in inflammatory responses.

Once we compared the patient data to the inflammatory signatures, we found three distinct groups of cancers: (1) those with high inflammation, (2) those with low inflammation, and (3) those with both high and low levels of inflammation. We found PAAD to be a member of the high inflammation group. This grouping is supported by multiple studies associating pancreatic inflammation (pancreatitis) with the development of pancreatic cancer (25, 32). One of the cancers we found to be in the low inflammation group was UVM. Melanomas are associated with environmental insults, such as exposure to ultraviolet light. As such, we expect that inflammation is not necessarily involved in the mechanism responsible for the development of skin cancer.

We believe that the gene expression data for these tumor types is heterogeneous across individuals, with multiple subgroups of patients per type. As such, the inflammatory signature presented in this first analysis is averaged across individuals, and that within these broad categories there may be subgroups of patients with high inflammation and others with low inflammation. This suggests that patients with these cancers could potentially benefit from further molecular subclassification.

In addition to correlating levels of inflammation with specific cancer types, we utilized clinical metadata from individuals with 7 different types of distinct tumors and associated this with molecular signatures derived from the web-based tool SaVanT. Molecular signatures are gene collections with associated biological interpretations that can identify genes upregulated in specific sample subsets compared to broader groups (15).

Signatures can be composed of genes associated with specific diseases. By performing a comparison of metadata with molecular signatures, we sought to evaluate if there was significant correlation between these values.

We found that four of the cancer types we evaluated (PAAD, CHOL, KICH, and UVM) had statistically significant associations between hallmark inflammatory response and at least one clinical variable. PAAD and KICH had a significant association with the patients’ stage of cancer at diagnosis, and CHOL and UVM had an association with vital status. Additionally, PAAD was significantly associated with sex. On average, females and individuals with stage II PAAD had the highest correlation between the clinical variable and hallmark inflammatory response. While for KICH, the highest average correlation was for individuals with stage IV cancer. Within each cancer type, alive individuals with CHOL and dead patients with UVM had the highest average correlation with hallmark inflammatory response. However, the correlation for both alive and dead vital status individuals was higher for CHOL than UVM.

Our results suggest that the use of molecular signatures in patients with cancer can provide valuable information. The signatures provided by SaVanT supplement MSigDB while utilizing the depth and specificity of large expression studies to describe the biology pertaining to various cancers and cell types. Having this information available for patients with cancer diagnoses could provide a deeper understanding of a patient’s clinical status. Furthermore, as there is marked heterogeneity even amongst specific organ-based tumors (Figure 2), molecular signatures could provide valuable information regarding the patient’s specific subtype of tumor.

In future studies we aim to further evaluate inflammation in cancer. Future studies could evaluate the relationship between additional tumor types with an expanded set of clinical variables. While we have shown three distinct groups of cancer types relative to inflammation levels, we also believe these results can be improved and expanded. For example, limiting the number of genes in the signatures or creating a cancer-specific inflammatory response panel of genes would produce a more cost-effective diagnostic test that could potentially be translated to the clinical setting.

In addition, although many cancer types fall into the high or low inflammation classifications, there are others with a mixed inflammation signal. The ambiguity in this group of cancer subtypes could arise from several sources and correcting for these sources may allow us to place these cancers into either the high- or low-inflammation group. For example, the mixed signal could be due to the need to subclassify patients even further for a particular cancer type. It is possible that some primary sites contain several populations of samples – such as those from a different biopsy type (i.e., blood or tumor sample). Determining these subgroups within the primary types would allow them to be treated independently.

Finally, gene expression data is one of many biological layers potentially contributing to cancer. Leveraging other levels of data, such as genome information, to identify mutations within tumors and patients would greatly assist in determining and developing therapies in a more personalized fashion for patients with different disease subtypes.

In summary, our study evaluated the association between inflammation signatures for different tumor types. We found associations between levels of inflammation and tumor types, and also found statistically significant relationships between patient metadata and inflammation for four tumor types. We believe our results demonstrate potential clinical utility in the continued establishment of personalized medicine and care for cancer patients, while further establishing the utility of SaVanT as a clinical tool.

